# Influence of Copper Dose on *Mycobacterium avium* and *Legionella pneumophila* Growth in Premise Plumbing

**DOI:** 10.1101/2025.03.20.644365

**Authors:** Rania E. Smeltz, Fernando A. Roman, Thomas Byrne, Rachel Finkelstein, Yang Song, Amy Pruden, Marc A. Edwards

## Abstract

Effects of copper at 0, 4, 30, 250, or 2000 µg/L on microbial communities were examined in an 11-month study using triplicate 120-mL water heater microcosms with PEX-b pipes containing mature biofilms to simulate premise plumbing. Effluent total cell counts (TCCs) and *Mycobacterium avium* peaked at 250 µg/L, reflecting the dual role of copper as a nutrient and antimicrobial. TCCs and *M. avium* were relatively consistent among replicate microcosms at each dose, but *Legionella pneumophila (Lp)* diverged among biological triplicates at 250 µg/L, consistently producing high culturable *Lp* (average 2.5 log MPN/mL) in one microcosm and low/non-detectable levels in the other two. Repeated cross-inoculations and a reinoculation failed to normalize the microbial community composition across 250 µg/L and other triplicate microcosms. 16S rRNA gene amplicon sequencing revealed that the 250 µg/L replicate with high *Lp* was characterized by a distinct microbial community composition relative to the two replicates. At 2,000 µg/L copper, microbial diversity and TCCs initially decreased, but then TCCs subsequently increased and ultimately were not statistically different from the 250 µg/L microcosms. This study provides insight into mechanisms underlying non-linear effects of copper dosing when applied as a disinfectant to premise plumbing for opportunistic pathogen control.

**SYNOPSIS:** Non-linear effects of copper on *Legionella pneumophila* and microbial communities are identified, providing deeper understanding to the efficacy of its use as a premise plumbing disinfectant.

## INTRODUCTION

Copper is of interest as a disinfectant for control of opportunistic pathogen growth in premise (i.e., building) plumbing. Copper can be released into drinking water from corrosion of copper alloy pipe or intentionally dosed for this purpose.^1^ Broadly, efficacy of copper as an antimicrobial can be dependent on water chemistry, physiology of the target microbes, biofilm characteristics, water flow patterns, levels of other disinfectants (e.g., chlorine), and other factors.^2,3^ Further, copper has been found to sometimes act as a nutrient to drinking water microbes at low concentrations and an antimicrobial at high concentrations,^2^ suggesting that sub-optimal doses could stimulate growth, rather than death, of pathogens. Thus, there is an array of factors to consider in applying copper as a premise plumbing disinfectant.

*Mycobacterium avium* and *Legionella pneumophila* (*Lp*) are two key opportunistic pathogens of concern that are prone to growth in premise plumbing.^2^ Prior field surveys of *Legionella* and *Lp* occurrence in drinking water suggest antimicrobial thresholds for copper of > 50 µg/L,^4^ > 400 µg/L,^5^ or > 1055 µg/L.^6^ However, other studies have reported contradictory results, suggesting positive effects on *Legionella* and *Lp* growth at concentrations > 500 μg/L^7^, or *Lp* persistence at concentrations > 2000 µg/L^8^. *M. avium* has been shown to be inactivated by high levels of copper in warm premise plumbing environments,^9,10^ but *M. avium* and other nontuberculous mycobacteria (NTM) are generally considered more resilient to copper than *Legionella*.^10,11^ The prior studies suggest that copper, if present at sufficiently high concentrations, could sometimes effectively control both NTM and *Lp,* which is the ideal case. Nonetheless, many existing strategies for controlling opportunistic pathogens in drinking water have limitations and, in some instances, may inadvertently promote pathogen proliferation.^11,12^ Controlled studies are needed to resolve discrepancies in the efficacy of copper as a disinfectant and to gain deeper mechanistic understanding needed to improve its application for simultaneous reduction of multiple opportunistic pathogens.

To gain insight into the effects of copper on *Lp* and NTM control in hot water plumbing systems, specifically at temperatures that support opportunistic pathogen growth in premise plumbing, we previously conducted a series of studies at both pilot- (spanning 3 years) and microcosm- (spanning 198 days) scale using the same local Blacksburg water supply.^9,13,14^ These studies demonstrated how a complex array of phenomena; including anode rod corrosion, associated hydrogen evolution, and pH shifts, can confound the antimicrobial action of copper. Similar confounding effects were likely at play, but not considered in the interpretation of results derived from prior field-scale research.^2^ Comparatively, another relatively long-term study (12 months) investigating opportunistic pathogens in water heaters was only able to detect NTM, and not *Legionella* spp., *Lp*, or *Pseudomonas aeruginosa* at any point in the experiment,^15^ highlighting challenges in simultaneous study of *Lp* and NTM at pilot scale, especially over extended time periods. In another prior microcosm-scale study, the effect of a 0.8 – 5 mg/L total Cu dose on *Lp* culturability was examined over a 4-week period using a synthetic water at 36°C.^16^ That work found that different *Lp* strains reacted differently to copper. Furthermore, environmental isolates derived from biofilm in a hot water system exhibited greater resistance to copper compared to the clinical strains tested, suggesting that the environmental isolates (which had a significantly higher expression of the copper resistance gene copA) were better adapted to copper.^16^ Finally, a 76 week microcosm-scale study examined the impact of different materials (including pipe coupons) on *Lp* and NTM, finding that high concentrations of copper (≈10 mg/L) suppressed established NTM and *Lp* and lowered their concentrations in the microcosms.^17^

The objective of this study was to assess the effects of a wide range of copper concentrations on *M. avium* and *Lp* in premise plumbing systems colonized by both organisms. Fifteen 120-mL water heater microcosms equipped with PEX-b pipes containing mature biofilms (>3.5 years old) and colonized with both *Lp* and *M. avium*^9^ were acclimated to Blacksburg tap water over a 6-month period and subsequently divided into biological triplicate microcosms subject to copper dosing at 0, 4, 30, 250, or 2,000 µg/L (as total copper) for 11-months. Microbial total cell counts (TCC) were monitored via flow cytometry, *M. avium* was measured via droplet digital polymerase chain reaction (ddPCR), and *Lp* was measured both via ddPCR and Legiolert™. 16S rRNA gene amplicon sequencing was applied to profile microbial communities and provide insight into microbial ecological factors mediating the effects of copper. The goal was to assess doses at which copper acted as a nutrient versus as an antimicrobial for opportunistic pathogens in warm water premise plumbing.

## MATERIALS AND METHODS

### Premise Plumbing Microcosm Design and Establishment

Fifteen new 120-mL glass microcosms (referred to as “simulated glass water heaters” in previous works)^18^ provided replicable simulation of premise plumbing with mature drinking water biofilm in triplicate. Each replicate microcosm contained two new and two conditioned PEX-b pipe coupons, with conditioned coupons cut from the middle of ∼4.6-meter recirculating lines of pilot-scale hot water plumbing systems, one that was previously dosed with phosphate and one that received no added phosphate during a >3.5-year-old experiment.^9,13^ All coupons were 2.5 cm long with a ∼ 1.7 cm inner diameter. The pilot-scale hot water plumbing systems had been inoculated with two strains of *Lp* serogroup-1 that were isolated from the Quincy, Illinois Veterans Home during a Legionnaires’ Disease outbreak.^19^ Prior to this experiment, the recirculation line biofilms were confirmed to be colonized by both *M. avium* and *Lp*,^9^ serving as an inoculum to the microcosms.

### Microcosm Influent & Water Changes: Acclimation Phase

The water heater microcosms were filled with 110 mL of local tap water (Blacksburg, VA) treated by the granular activated carbon (GAC) filter feeding the recirculating pipe rig from which the pipe coupons were extracted. The influent water was collected in batches at 5-week intervals and stored at 4°C. On the day before a water change, 2L of the water was collected and heated to 37°C overnight then adjusted to pH 6.70 ± 0.05 using H_2_SO_4_, before addition to the microcosms. The microcosm bulk water was changed 2× weekly, by replacing 75% of the bulk water volume (∼82 mL) in each microcosm with new influent, to match the 3.2-day hydraulic retention time of the pilot-scale plumbing system. The 3.2-day hydraulic retention time was selected to simulate common residential hot water usage in the United States.^13^ The microcosms were maintained at 37°C, representing the low end of a water heater set point, and gently mixed on a shaker table at 100 rpm. The microcosms were acclimated for 6 months under these conditions prior to commencement of copper dosing.

### Cross-Inoculation of the Microcosms

A month into the acclimation phase, the effluent from each microcosm was cross inoculated with the aim of establishing baseline *Lp* and microbial populations across the 15 replicates. This was achieved by collecting the effluent (75% bulk water volume) from the microcosms into a common reservoir (an autoclaved 2-L polypropylene bottle), vigorously shaking this mixture, returning 25% of the combined effluent to the microcosms, and replacing the remaining 50% with the GAC-treated water. This procedure was repeated over the course of ten sequential water changes, starting a month into the acclimation phase, for five weeks thereafter.

### Copper Dosing and Reinoculation

Nearing the end of the 6-month acclimation phase, a week before the copper dosing commenced, the microcosms were sampled for culturable *Lp* levels measured by Legiolert^TM^ tests (IDEXX Laboratories, Westbrook, ME) and TCCs. Based on these measurements, the microcosms were grouped into five sets of triplicate microcosms, so that there was no statistical difference in mean values of TCC or *Lp* among experimental conditions (ANOVA, p > 0.05) (SI Figure 1). Triplicate microcosms subsequently received one of five copper dosages of 0, 4, 30, 250, or 2000 µg/L copper over an 11-month period. Copper was dosed directly to the influent using stock solutions prepared with CuSO_4_ ·5H_2_O. After ∼ 3.5 months of copper dosing, random testing among the microcosms indicated that *Lp* still had not established in some replicates and, therefore, all microcosms were reinoculated once more. This was achieved using water from the same pilot-scale water heater from which the microcosm pipe coupons were originally derived. The inoculation water was mixed with the normal influent water to target ∼50 MPN/mL culturable *Lp* in each microcosm.

**Figure 1.**
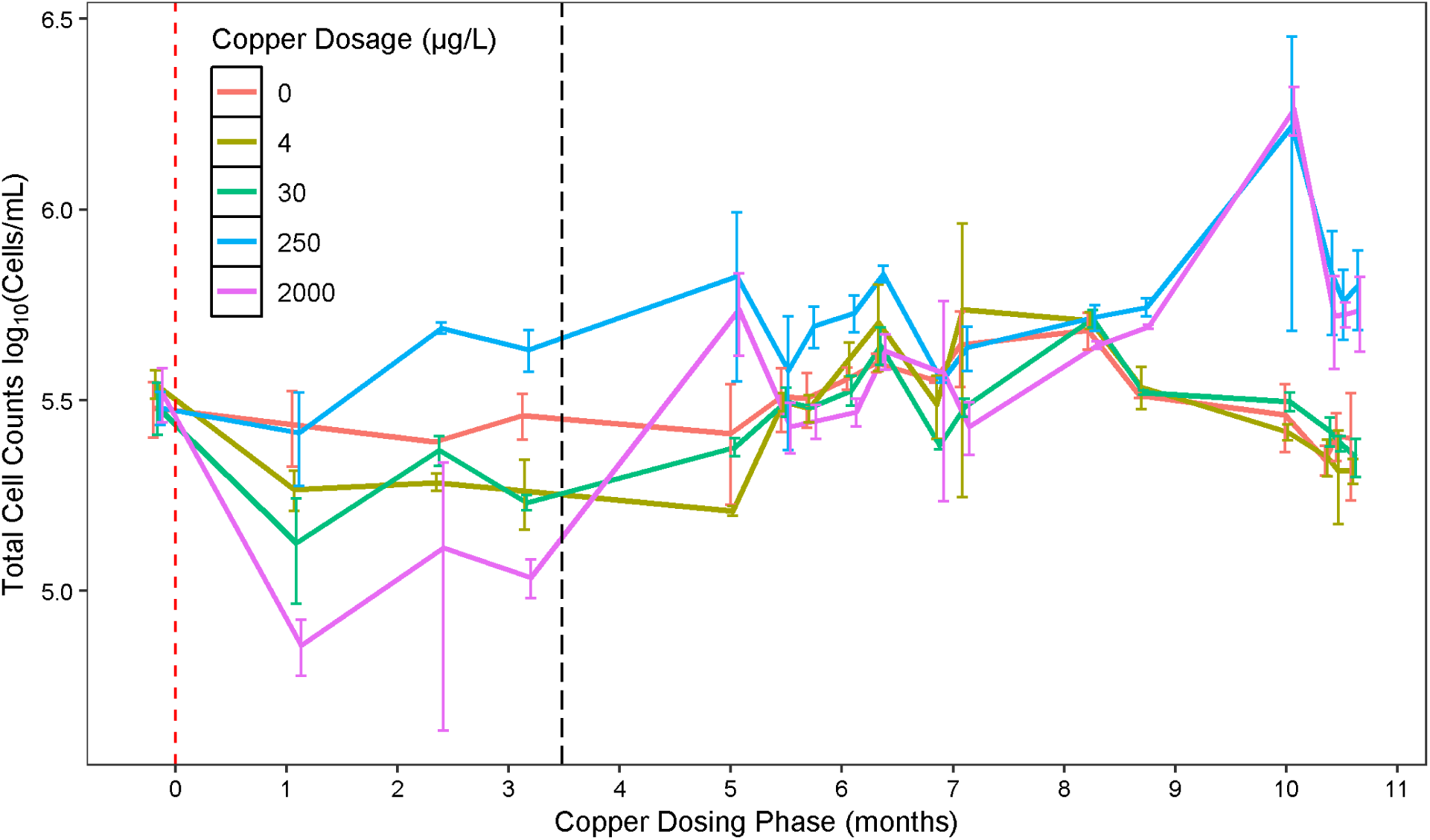
Mean Total Cell Counts (cells/mL) in microcosm bulk water effluent over the course of the experiment. Error bars are the standard deviation of biological triplicate microcosms. Lines connect the data points to guide comparison and are colored by copper concentration. The first measurement (time −0.25) occurred at the end of the acclimation phase, a week before copper dosing began (time = 0 represented by the red dashed vertical line). The black dashed vertical line at ∼ 3.5 months represents the point at which all reactors were reinoculated with *Lp*. Time points have a set jitter to avoid overlapping datapoints.

The influent water preparation protocol was changed throughout the 6-month acclimation (experimental months 0-6) and 11-month copper dosing (experimental months 6-17) phases of the experiment in an attempt to establish a replicable response of *Lp* to copper. Modifications included: 1) passing GAC-filtered water through a ferric oxide filter to decrease phosphate levels to < 0.05 mg/L followed by filter-sterilizing the water using a 0.22-μm pore size mixed nitrocellulose ester membrane (Whatman, Maidstone, United Kingdom) during the experimental months 0-8; 2) dosing GAC-treated influent with phosphate throughout the copper dosing phase to achieve 5 μg/L as P during experimental months 6-17; 3) no longer sterilizing the influent or treating it with a ferric oxide filter during experimental months 8-17; and 4) dosing ferric pyrophosphate (100 µg as Fe) after week 35 of the copper dosing phase to provide bioavailable iron experimental months 14-17.

### Microbial Analysis

Effluent collected from the microcosms during bi-weekly water changes was subject to microbial analysis. TCC was measured on a BD Accuri C6 (BD Bioscience, Franklin Lakes, NJ) using SYBR Green I dye to stain total (intact + damaged) cells.^20,21,22^ Gating used the Eawag FL1-A (emission filter 533/30) vs FL3-A (emission filter 670 LP) template for drinking water, with 50 μL of each sample analyzed at medium speed (35 μL/min) and with an acquisition threshold set to 800.^22^ Legiolert^TM^ was used to enumerate culturable *Lp*, subjecting 1.0 mL of sample bulk water to the nonpotable procedure per the manufacturer’s instructions.

During regular water changes, ∼82 mL of influent or effluent water from each microcosm was filtered through a 0.22-µm mixed-cellulose ester membrane filter (MilliporeSigma, Burlington, MA) and the filter was subject to DNA extraction using a FastDNA SPIN kit (MP Biomedicals Inc., Solon, OH).

### Droplet Digital PCR (ddPCR) Analysis

DNA extracts were analyzed for opportunistic pathogens using a QX200 ddPCR instrument (Bio-Rad, Hercules, CA) (SI Table 1). These assays, targeting the *Lp* specific *mip* gene and *M. avium* 16S rRNA gene, had previously been optimized for qPCR^23^ and were adapted for ddPCR in this study. Thermal gradients, ranging from 50-60°C, were conducted on each assay to determine the optimal annealing temperature (SI Table 1) to separate negative and positive droplets. Each sample was analyzed in technical triplicate for *Lp* and *M. avium.* Reactions were prepared using a total volume of 22 µL per well, with 20 µL in each well used for analysis. PCR amplification was carried out on a C100 Touch Thermal Cycler (Bio-Rad) (thermocycling conditions listed in SI Table 1). Each 96-well plate included three no-template controls (NTCs) using molecular-grade water, as well as three positive controls containing synthetic gene fragments (gBlocks™, Integrated DNA Technologies, Coralville, IA) corresponding to the targeted gene regions. A sample was classified as positive if a minimum of three positive droplets were detected in at least two of the three of the technical replicates.^24^ Only data from wells generating > 10,000 accepted droplets were included in the analysis.

### 16S rRNA Gene Amplicon Sequencing and Data Analysis

To analyze the microbial composition of each microcosm, 16S rRNA gene amplicon sequencing was carried out on the DNA extracts. Amplicon sequence reads targeted the V4-V5 hypervariable region of the 16S rRNA gene using the 515f/926r primer set. Samples were sequenced on a MiSeq V3 600 cycle run, yielding an average of 119,910 reads per sample (SD = 69,156). Reads were imported and subsequent analysis were carried out using DADA2 (v1.18.0)^25^ in R (v 4.0.3).^26^ Reads were filtered, trimmed, and paired before exact amplicon sequence variants (ASVs) were generated and chimeras removed. Taxonomy of ASVs was assigned using RDP Train Set 18.^27^ ASVs were rarefied to the minimum sequencing depth of 10,544.^28^ Multivariate homogeneity was checked with the betadisper R function (p > 0.05) before conducting permutational multivariate analysis of variance (PERMANOVA) with Bray-Curtis distance matrices using the vegan R package (v 2.6.4).^29^ Linear mixed-effect models were created on *M. avium* and *Lp* ddPCR measurements, controlling for the three consecutive samplings of each microcosm in the final month of the experiment using the lme4 R package (v 1.1-36).^30^ ANOVA was run on the linear mixed-effect models to assess the impact of copper dosage and on the Shannon and Simpson diversity (calculated with vegan) using the R stats package (v 4.4.2).^26^ Tukey’s HSD was used as a post-hoc test to ANOVA to assess the impact of each copper dosage using the multcomp R package (v 1.4-28).^31^ Wald’s test was used for the log2 fold change between the 250 µg/L Cu condition microcosms using the DESeq2 R package (v 1.46.0).^32^A paired t-test controlling for repeated measures was used to assess differences in TCC before and after copper dosing using the R stats package (v 4.4.2).^26^

### Data Availability

The 16S rRNA gene amplicon sequencing gathered from the microcosms in this experiment has been uploaded to the National Center for Biotechnology Information Sequence Read Archive database under Accession number PRJNA1229645.

## RESULTS

### Total Cell Counts

The effluent mean TCC for all the microcosms during the end of the acclimation phase (month 6), before copper addition, was 3.18×10^5^ cells/mL (SD = 4.16×10^3^) (Figure 1, SI Figure 1b). In the short-term after copper dosing began (1-3.5 months post copper dosing), the TCC indicated growth at 250 µg/L. As the experiment progressed, the three replicates dosed with 250 µg/L copper consistently produced the highest cell counts. Replicates dosed with 2000 µg/L copper initially experienced a 0.66 log reduction (a 78% decrease from its final pre-dosing concentration), resulting in the lowest TCCs observed in the study. But by the end of the 11-month experiment, the mean TCC at 2000 µg/L copper gradually recovered and eventually surpassed those at doses ≤ 30 µg/L and were not significantly different than at 250 µg/L (Tukey’s HSD p<.05) (Figure 1, SI Table 2). In fact, the TCC steadily increased with time in the 2000 µg/L microcosms, over a time period spanning both before and after the re-inoculation attempt that occurred 3.5 months into the copper dosing phase (Figure 1, SI Figure 2).

### *Mycobacterium avium* response to copper in the microcosms

When tested over three sequential samplings in the final (11^th^) month of the experiment, significant differences in *M. avium* gene copies/mL were found in the microcosm bulk water as a function of copper concentration (Linear Mixed-Effect Model ANOVA, p<0.05) (Figure 2). The 250 µg/L condition contained higher concentrations of *M. avium* compared to other copper dosages (0.55 – 0.85 logs higher) (Tukey’s HSD, p<0.05, SI Table 3). *M. avium* was not detectable in the influent water using ddPCR, indicating that the increase in the effluent was the result of growth in the microcosms.

**Figure 2.**
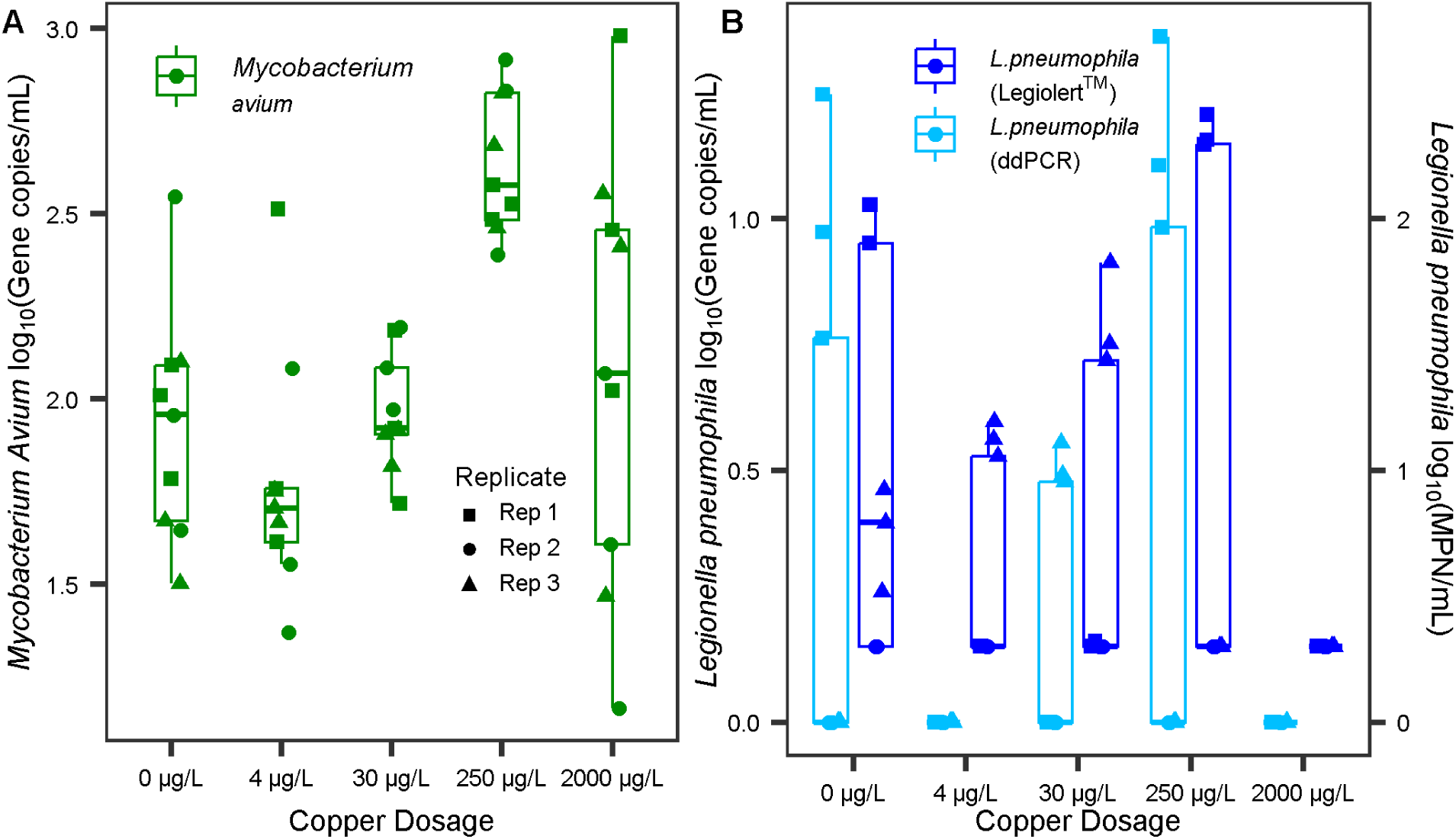
Mean ddPCR technical triplicate gene copies/mL counts of (A) *M. avium* detected through ddPCR and (B) *Lp* detected via ddPCR (mean technical triplicate) and Legiolert™ across the fifteen microcosms over three consecutive samplings in the final (11^th^) month of the experiment. For (A), the data consists of 5 copper levels × 3 microcosms × 3 sampling events = 45 *M. avium* data points). For (B) the data consists of 5 copper levels × 3 microcosms × 3 sampling events = 45 culturable (MPN/mL) or 45 ddPCR (gc/mL) *Lp* data points. *M. avium* was detectable in all samples. For *Lp*, points plotted at zero (gc/mL or MPN/mL) indicate non-detects for ddPCR (estimated ∼detection limit of 1.5 gc/mL) and Legiolert^TM^ (detection limit of 1 MPN/mL for the protocol used).

### *L. pneumophila* response to copper in the microcosms

*Lp* response to copper dosing was confirmed both by ddPCR and Legiolert^TM^. On dates when both Legiolert^TM^ and ddPCR tests were conducted on aliquots of the same sample (n=45), measurements were strongly correlated (R^2^ = 0.85) (t-test, p<0.05), but the log ddPCR gc/mL values were typically 0.51 times the log of Legiolert^TM^ MPN/mL (SI Figure 3). *Lp* concentrations did not significantly differ across the microcosms as a function of copper condition (Linear Mixed-Effect Model ANOVA, p > 0.05), but both Legiolert^TM^ and ddPCR measurements indicated a consistent ranking of average concentration across microcosms as follows: 2000 µg/L ≤ 4 µg/L < 30 µg/L < 0 µg/L < 250 µg/L. *Lp* was never detectable in the influent used for the microcosm water changes using ddPCR or Legiolert^TM^, indicating that the increase in the effluent was the result of growth in the microcosms. It was noted that there was a much wider variance in *Lp* numbers among replicates than there were for *M. avium* (Figure 2). This was most evident for microcosms dosed with 250 µg/L copper, for which one microcosm consistently displayed the highest concentrations of culturable *Lp*, while the second and third replicates produced non-detectable levels of culturable *Lp* over the majority of the sampling events (87% and 73% of events) over the course of the copper-dosing phase of the experiment (Figure 3). *Lp* levels quantified via Legiolert^TM^ ranged from 8 MPN/mL to 1,460 MPN/mL for the first replicate microcosm in the 250 µg/L Cu condition (Figure 3). The stark differences between these microcosms were maintained even though they had been cross-inoculated ten times during acclimation to establish the same baseline microbial community prior to copper dosing, reinoculated ∼ 3.5 months into the copper dosing phase, and subject to four adjustments to the water chemistry.

**Figure 3.**
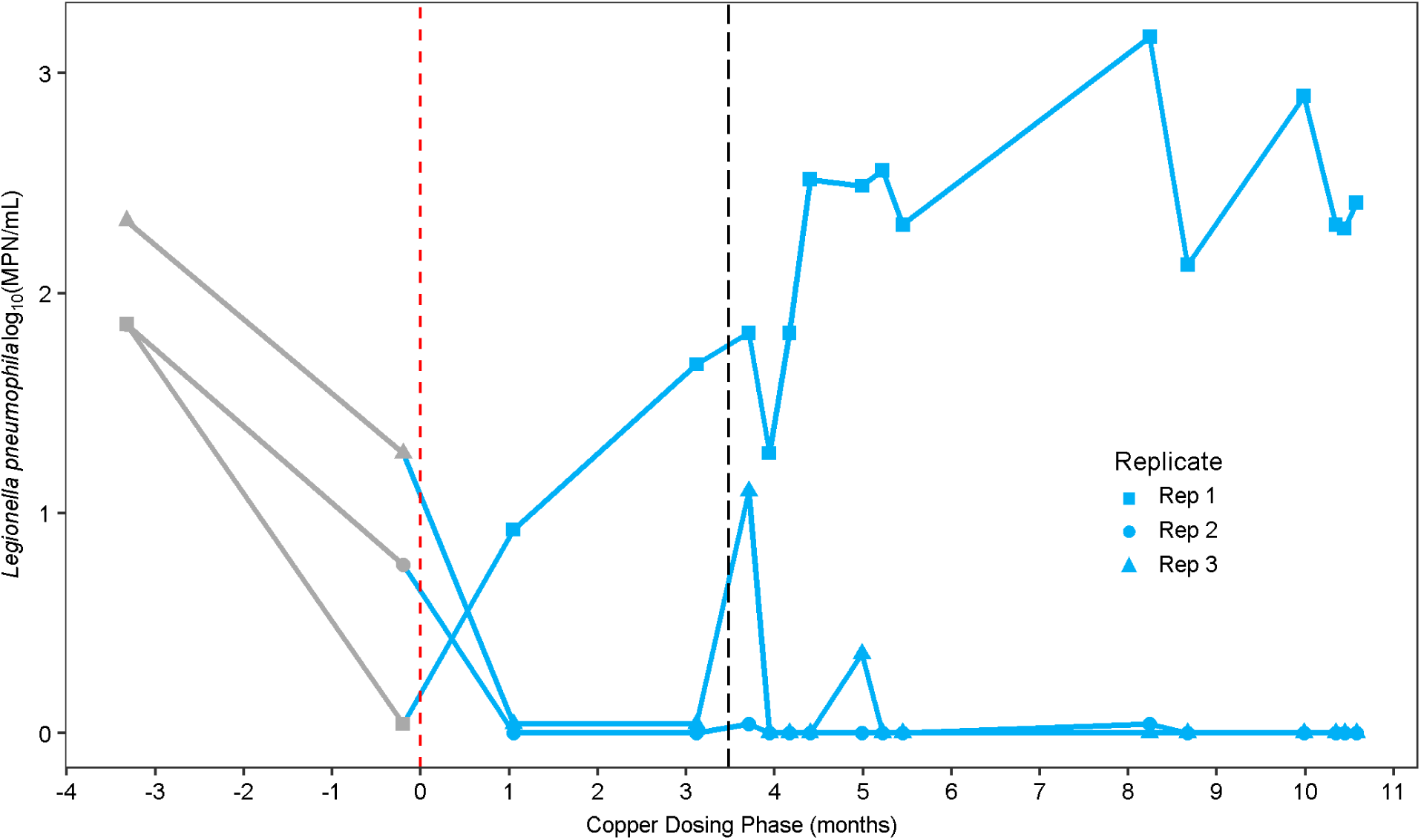
*Lp* MPN/mL, determined via Legiolert™, among the three replicate microcosms dosed with 250 µg/L copper that displayed divergent behavior over the 11-month experimental period. The vertical, dashed line at ∼3.5 months represents the point at which all microcosms were reinoculated with *Lp*. Time 0, shown with the red dashed vertical line represents when the copper dosing began. Points before 0 months represent culturable *Lp* in the microcosms prior to copper dosing, during the acclimation phase. Culturable *Lp* concentration measured with time for the microcosms receiving other copper concentrations can be found in SI Figure 4. Legiolert™ detection limit was 1 MPN/mL. Points plotted at an MPN/mL of zero indicate non-detects for culturable *Lp*.

Consistent with the Legiolert^TM^ results, replicates 2 and 3 of the 250 µg/L copper condition consistently yielded undetectable gene copies/mL of *Lp*, while up to 22 gene copies/mL were measured in replicate 1 (Figure 2).

### 16S rRNA gene amplicon Sequencing

16S rRNA gene amplicon sequencing allowed comparison of the microbial community profiles across the microcosms and influent water (Figure 4). The microbial communities were significantly different from each other as a function of copper concentration (PERMANOVA, p < 0.05) (Figure 5). Copper doses of 0, 4, and 30 µg/L produced a similar microbial profile in terms of the ten most abundant phyla from each replicate microcosm, with a marked shift in composition in the 250 µg/L and 2,000 µg/L conditions (Figure 4). Of note, the microcosms dosed with 0, 4, and 30 µg/L copper were mostly comprised of *Proteobacteria*, while the reactors dosed with 250 µg/L copper contained relatively equal proportions of *Proteobacteria* and *Gemmatimonadetes.* Microcosms dosed with 2000 µg/L were primarily dominated by *Bacteriodetes* and *Proteobacteria* (Figure 4). Within each condition from 0-250 µg/L, the microbial communities appeared relatively consistent among the three replicates. However, there was marked variation in the microbial community among the 2,000 µg/L condition replicates (Figure 5). Copper concentration had a significant effect on Shannon diversity and Simpson diversity (ANOVA, p < 0.05) (Figure 6).

**Figure 4.**
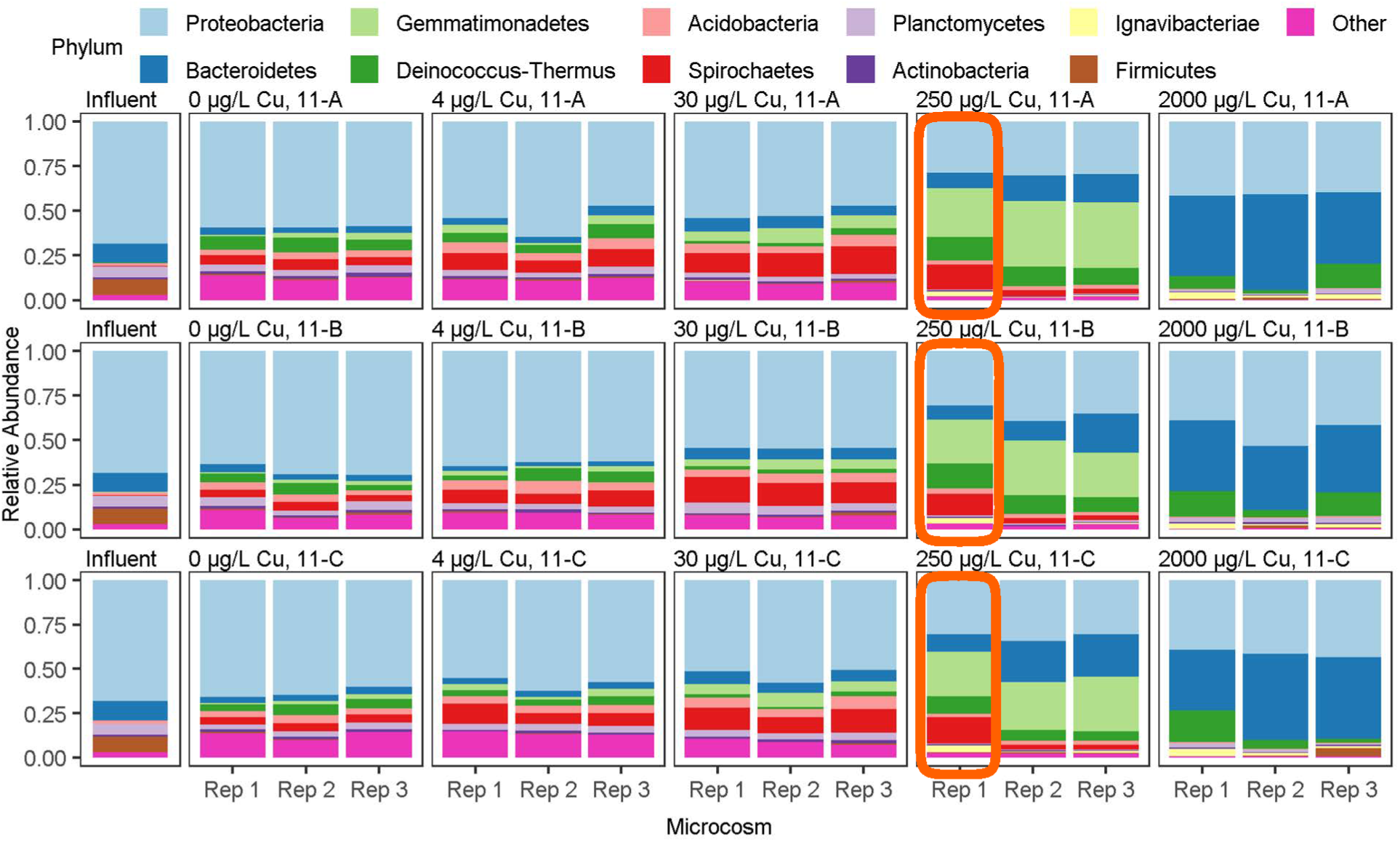
Relative abundances of each of the top ten most abundant phyla over all ASVs in each replicate microcosm grouped by copper concentration and sampling date. The orange boxes indicate the 250 µg/L copper replicate that consistently displayed the highest concentrations of *Lp*. The influent [collected at the start of month 10 for water changes (n=1)] represents the water used throughout three sequential sampling events during water changes carried out in the final (11^th^) month of the experiment. 11-A/B/C, represent the first (A), second (B), and third (C) sequential water changes.

**Figure 5.**
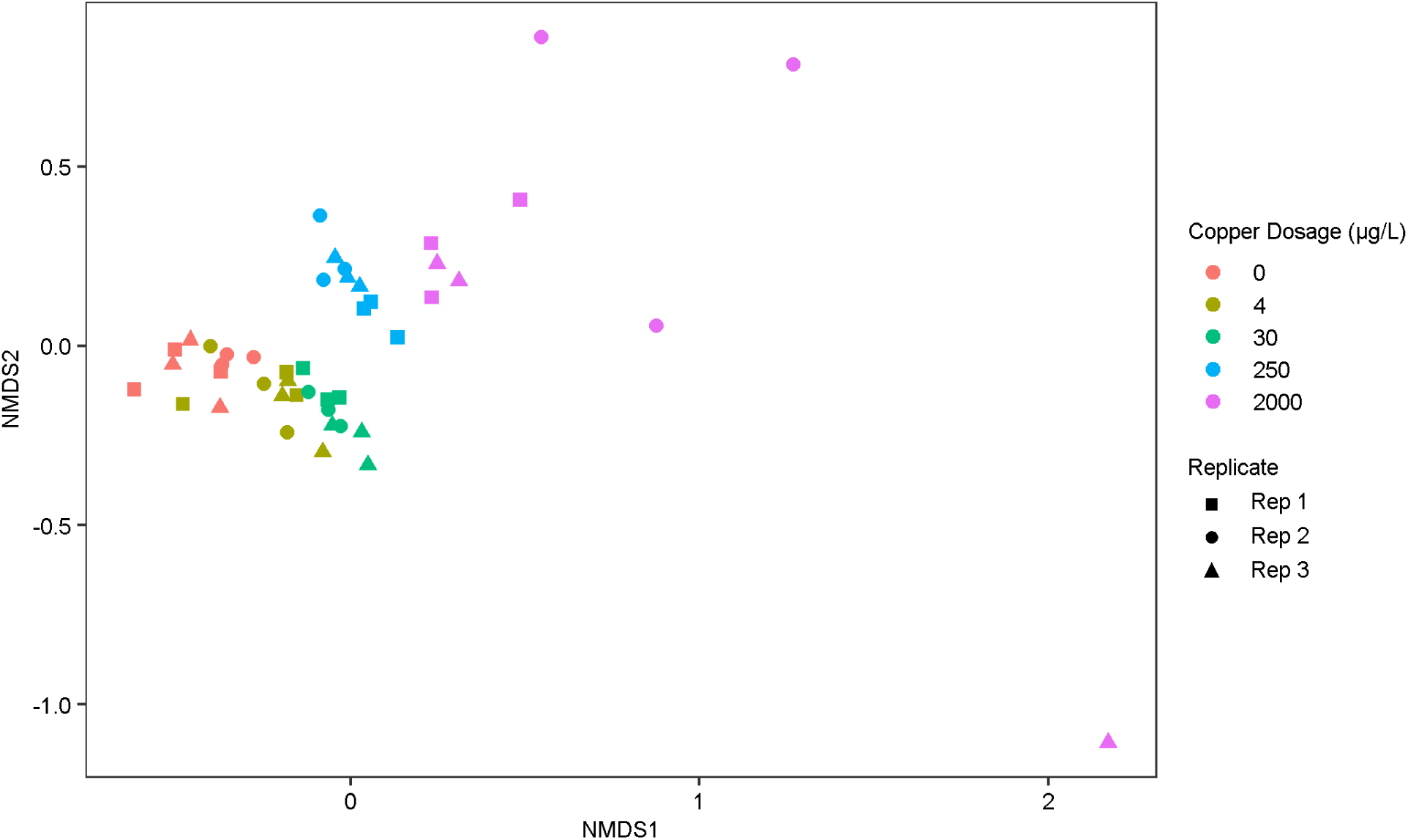
Bray-Curtis-based nonmetric multidimensional scaling of taxa from effluent water of microcosms dosed with 0, 4, 30, 250, and 2000 µg/L of copper over an 11-month period (Stress = 0.08836). The data consists of 5 copper levels × 3 microcosms (per copper level) × 3 sampling events = 45 data points.

**Figure 6.**
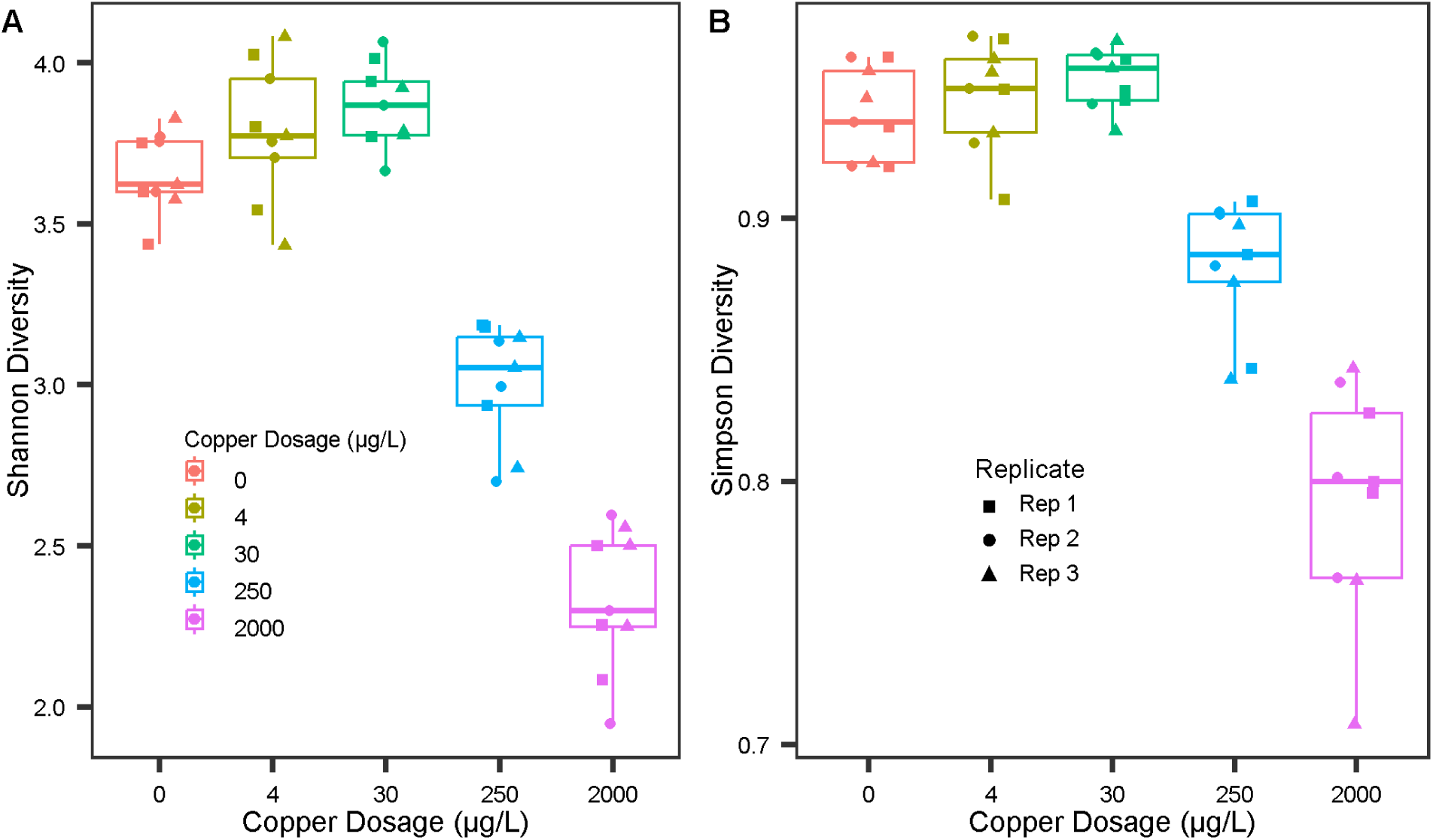
Shannon and Simpson diversity index of replicate microcosms on the final three sampling dates grouped by copper condition. The data consists of 5 copper levels × 3 microcosms (per copper level) × 3 sampling events = 45 data points.

The microbial community profile of the 250 µg/L microcosm that consistently had the highest level of *Lp* was found to be dominated by the same top ten phyla, but differences in relative abundance stood out (Figure 4, highlighted in orange boxes). Specifically, this microcosm was significantly enriched in *Ignavibacteriae* and depleted in *Chlamydiae* and *Bacteroidetes* (Wald’s test, p<0.05) (Figure 4, highlighted in orange boxes). Beta diversity analysis, however, did not reveal an obvious difference in the microbial community composition of this replicate beyond the variation that was seen for replicates of other copper concentrations (Figure 5).

Relative abundances of individual classes of bacteria were further compared across microcosm conditions, revealing some notable differences as a function of copper concentration (Figure 7). For example, *Proteobacteria* was higher in the 2000 µg/L than in the 250 µg/L condition, while *Firmicutes* indicated little change across copper dosages (Figure 7). *Gemmatimonadetes* were enriched from 0 to 250 µg/L, but dramatically decreased at 2000 µg/L (Figure 7). *Deinococcus-Thermus* exhibited a slight decrease in relative abundance from 250 µg/L to 2000 µg/L (Figure 7). *Chloroflexi*, *Acidobacteria*, and *Actinobacteria* markedly decreased with increasing copper (Figure 7).

**Figure 7.**
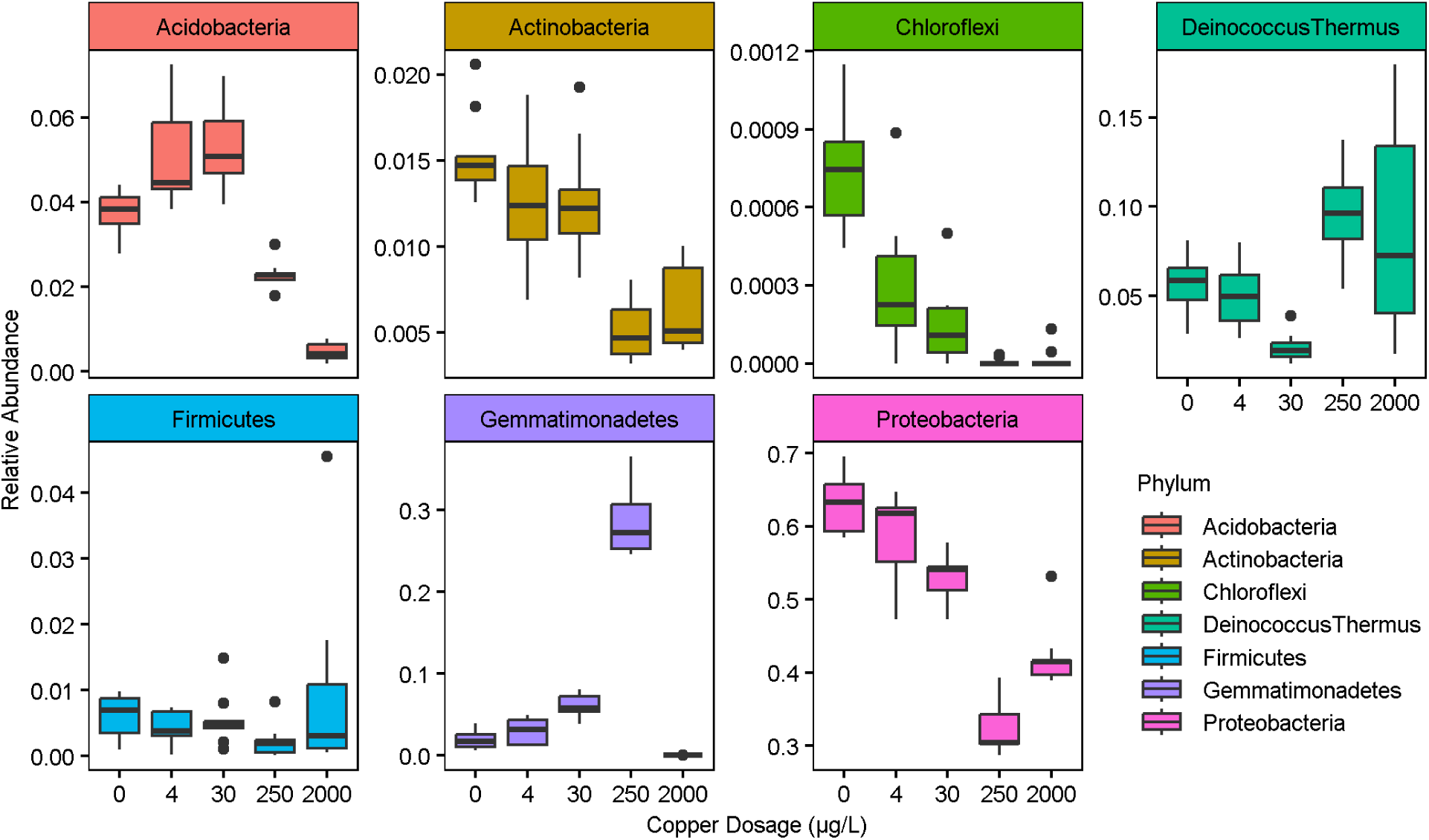
Relative abundance of select phyla across copper dosages in each replicate over the three sequential sampling events carried out in the final (11^th^) month of the experiment.

## DISCUSSION

This study reveals apparent hormetic effects of copper on premise plumbing microbiomes, i.e., acting as a nutrient at low concentrations and as an antimicrobial at higher concentrations. TCCs provided an indicator of the effects on bacterial communities at-large, revealing some surprising patterns over the course of this near year-long experiment. While TCCs initially decreased markedly when the highest dose of 2000 µg/L was commenced, as one would expect, over the course of subsequent months, the microbial community adapted such that TCCs were as high as the 250 µg/L by the end of the experiment. Further, TCC levels in the 2000 µg/L condition had not yet plateaued by the end of the experiment, suggesting that levels could have eventually surpassed those of the 250 µg/L condition.

Interestingly, TCCs initially increased in the 250 µg/L condition and remained elevated throughout the experiment relative to the un-dosed control, suggesting that copper acted largely as a nutrient at this concentration. Doses of 4 and 30 µg/L Cu were initially associated with a significant decrease in TCCs (paired t-test, p < 0.05), but remained comparable to the control for the majority of the experiment.

*M. avium* concentrations, which were only measured in the 11^th^ month, were also highest in the 250 µg/L condition, but were lowest in the 2000 µg/L condition. This suggests that very high doses of copper are likely needed to effectively control *M. avium*, and lower doses serve as a nutrient or selector for the organism. Overall, the findings for TCC and *M. avium* were consistent with expectations that copper has a dual role of nutrient and antimicrobial.^2^ Further, a level of copper that is antimicrobial in the short-term can behave as a nutrient over a period of months to years as microbial communities adapt to the copper.^16^

Support for this hypothesis was provided in the microbial community analysis. Even though TCCs were elevated in the 250 µg/L and 2000 µg/L conditions, microbial diversity was diminished. As richness and evenness declined, a subset of the remaining species were enriched and became dominant in the community (Figures 4 and 6). With the significant decrease in diversity, a few taxa prevailed, making the community vulnerable to stochasticity. This was apparent in terms of the wide variance in microbial community composition among the 2000 µg/L replicates (Figure 5).

Prior research by Song and colleagues using the same municipal tap water supply also assessed the effect of dosed copper on the microbial community in a full-scale water system, but the experimental design differed from the present study in that the dose was incrementally increased from 0, 50, 100, 300, 600, 1200, to 2000 µg/L, while maintaining each concentration for only periods of 4 to 16 weeks.^9^ Overall Phyla-level trends in *Proteobacteria*, *Deinococcus-Thermus*, *Chloroflexi*, *Acidobacteria*, and *Actinobacteria* were similar to those observed in this work. However, Song and colleagues also observed enrichment over 6 weeks at 1200 µg/L in *Firmicutes*, and enrichment of *Gemmatimonadetes* up through 2000 µg/L. The differences in *Firmicutes* and *Gemmatimonadetes* behavior when comparing the results of this study could potentially be attributed to differences in duration of copper dosing, as Song and colleagues maintained the 2000 µg/L copper dose for 12 weeks,^9^ whereas the microcosms in this study were acclimated at this dose for 11 months.

At 250 µg/L, stochasticity was observed in the concentrations of culturable *Lp*, measurements of which maintained relatively consistent among all replicates at lower copper doses (0, 4, and 30 µg/L total Cu), while almost always being undetectable at 2000 µg/L (SI Figure 4). In a prior study investigating the effect of copper pipe in warm premise plumbing systems, *Lp* numbers were initially lowest in the biofilms on copper pipes, but eventually increased to the same concentrations as PEX and stainless-steel pipes after two years.^33^ In such studies, it is important to note that copper pipe aging leads to decreasing release of copper into solution, and thus it is difficult to know if the observed increase in *Lp* was due to lower levels of copper or microbial adaptation.

A synthesis of our results over the last decade using Blacksburg tap water reveals two general responses of experimental plumbing systems with respect to *Lp* inoculation. As one would expect, there are many circumstances in which the system response to inoculated *Lp* is deterministic, in which replicate microcosms behave as true replicates. But here, we report a situation in which stochastic behavior clearly occurred, because in some replicate microcosms *Lp* consistently died off and in other reactors it thrived. With benefit of hindsight, we now recognize similar stochastic behavior for *Lp* amongst replicate microcosms during at least two prior experiments in our lab.^34,35^ We were reluctant to accept the conclusion that these systems were stochastic, preferring to believe that there were subtle uncontrolled differences between replicates, but we consider the evidence gathered herein to be conclusive. This study demonstrates that even in the most simplistic simulation of premise plumbing, i.e., a glass microcosm containing pipe sections, that there are complexities at play that can determine if effluent *Lp* is extremely high or very low.

Interestingly, the replicate microcosms described herein were deterministic and replicable for measured dimensions of cell counts, mycobacteria and community analysis, even as they were stochastic for *Lp*. We speculate that the sensitive predator, prey and parasitic relationships that drive the *Lp* life cycle make it particularly vulnerable to random events that can dictate the trajectory of its ecology in the premise plumbing environment. In a companion paper we report a discovery that *Neochlamydia* was consistently abundant in the replicate microcosms where *Lp* failed to establish.^36^ Previous research in pure culture settings has shown that amoeba carrying *Neochlamydia* are resistant to *L. pneumophila* infection,^37^ potentially explaining the observed stochastic behavior. If such stochastic behavior for *Lp* is commonplace, as it is in certain other areas of microbial ecology,^38,39^ this discovery has profound implications for experimental design and the interpretation of past data from laboratory and field sampling.^40^ Overall, the findings of this study reveal both hormesis and stochasticity in microbial ecological succession as key mechanisms underlying the observed non-linear response of *Lp* to copper as a disinfectant in premise plumbing.

## ASSOCIATED CONTENT

### Supporting Information

Information regarding ddPCR assays, statistical analysis, initial *Lp* and TCC of the microcosm groups at the end of the acclimation phase, additional TCC information for the 2000 µg/L Cu condition, a comparison between Legiolert^TM^ and ddPCR *Lp* measurements, and the culturable *Lp* for all the copper conditions throughout the study, can be found in the Supporting Information document (PDF).

### Author Contributions

R.E.S, F.A.R., T.B., and R.F. are designated as co-first authors and contributed equally to this work.

### NOTES

The authors declare no conflict of interest.

## ACKNOWLEDGEMENTS

This study was primarily funded using Dr. Edwards’ and Dr. Pruden’s discretionary funding. In addition, this work was supported in part by the National Science Foundation Award 2125798. Any opinions, findings, and conclusions or recommendations expressed in this material are those of the author(s) and do not necessarily reflect the views of the National Science Foundation.

## REFERENCES

1. LeChevallier, M. Examining the Efficacy of Copper-Silver Ionization for Management of Legionella: Recommendations for Optimal Use. AWWA Water Science 2023, 5 (2). DOI:10.1002/aws2.1327.

2. Cullom, A. C.; Martin, R. L.; Song, Y.; Williams, K.; Williams, A.; Pruden, A.; Edwards, M. A. Critical Review: Propensity of Premise Plumbing Pipe Materials to Enhance or Diminish Growth of Legionella and Other Opportunistic Pathogens. Pathogens 2020, 9 (11), 957. DOI:10.3390/pathogens9110957.

3. Zhang, Y. Nitrification in Premise Plumbing and Its Effect on Corrosion and Water Quality Degradation; Ph.D. Dissertation, Virginia Tech, 2008. https://vtechworks.lib.vt.edu/items/d57ec849-d055-4852-b34e-744a636f46ce (accessed 2025-08-01).

4. Bargellini, A.; Marchesi, I.; Righi, E.; Ferrari, A.; Cencetti, S.; Borella, P.; Rovesti, S. Parameters Predictive of Legionella Contamination in Hot Water Systems: Association with Trace Elements and Heterotrophic Plate Counts. Water Research 2011, 45 (6), 2315–2321. DOI:10.1016/j.watres.2011.01.009.

5. Liu, Z.; Stout, J. E.; Tedesco, L.; Boldin, M.; Hwang, C.; Diven, W. F.; Yu, V. L. Controlled Evaluation of Copper-Silver Ionization in Eradicating Legionella Pneumophila from a Hospital Water Distribution System. Journal of Infectious Diseases 1994, 169 (4), 919–922. DOI:10.1093/infdis/169.4.919.

6. Miuetzner, S.; Schwille, R. C.; Farley, A.; Wald, E. R.; Ge, J. H.; States, S. J.; Libert, T.; Wadowsky, R. M. Efficacy of Thermal Treatment and Copper-Silver Ionization for Controlling Legionella Pneumophila in High-Volume Hot Water Plumbing Systems in Hospitals. American Journal of Infection Control 1997, 25 (6), 452–457. DOI:10.1016/s0196-6553(97)90066-3.

7. Mathys, W.; Stanke, J.; Harmuth, M.; Junge-Mathys, E. Occurrence of Legionella in Hot Water Systems of Single-Family Residences in Suburbs of Two German Cities with Special Reference to Solar and District Heating. International Journal of Hygiene and Environmental Health 2008, 211 (1–2), 179–185. DOI:10.1016/j.ijheh.2007.02.004.

8. Mathys, W.; Palma Hohmann, C.; Junge-Mathys, E. Efficacy of Copper-Silver Ionization in Controlling Legionella in a Hospital Hot Water Distribution System: A German Experience. Wiley Online Library 2001. DOI:10.1128/9781555817985.ch84.

9. Song, Y.; Finkelstein, R.; Rhoads, W.; Edwards, M. A.; Pruden, A. Shotgun Metagenomics Reveals Impacts of Copper and Water Heater Anodes on Pathogens and Microbiomes in Hot Water Plumbing Systems. Environmental Science & Technology 2023, 57 (36), 13612–13624. DOI:10.1021/acs.est.3c03568.

10. Lin, Y. E.; Vidic, R. D.; Stout, J. E.; McCartney, C. A.; Yu, V. L. Inactivation of Mycobacterium Avium by Copper and Silver Ions. Water Research 1998, 32 (7), 1997–2000. DOI:10.1016/s0043-1354(97)00460-0.

11. Kusnetsov, J.; Iivanainen, E.; Elomaa, N.; Zacheus, O.; Martikainen, P. J. Copper and Silver Ions More Effective against Legionellae than against Mycobacteria in a Hospital Warm Water System. Water Research 2001, 35 (17), 4217–4225. DOI:10.1016/s0043-1354(01)00124-5.

12. Donohue, M. J.; Vesper, S.; Mistry, J.; Donohue, J. M. Impact of Chlorine and Chloramine on the Detection and Quantification of Legionella Pneumophila and Mycobacterium Species. Applied and Environmental Microbiology 2019, 85 (24). DOI:10.1128/aem.01942-19.

13. Song, Y.; Pruden, A.; Rhoads, W. J.; Edwards, M. A. Pilot-Scale Assessment Reveals Effects of Anode Type and Orthophosphate in Governing Antimicrobial Capacity of Copper for Legionella Pneumophila Control. Water Research 2023, 242, 120178. DOI:10.1016/j.watres.2023.120178.

14. Rhoads, W. J.; Pruden, A.; Edwards, M. A. Interactive Effects of Corrosion, Copper, and Chloramines on Legionella and Mycobacteria in Hot Water Plumbing. Environmental Science & Technology 2017, 51 (12), 7065–7075. DOI:10.1021/acs.est.6b05616.

15. Tolofari, D. L.; Bartrand, T.; Masters, S. V.; Duarte Batista, M.; Haas, C. N.; Olson, M.; Gurian, P. L. Influence of Hot Water Temperature and Use Patterns on Microbial Water Quality in Building Plumbing Systems. Environmental Engineering Science 2022, 39 (4), 309–319. DOI:10.1089/ees.2021.0272.

16. Bédard, E.; Trigui, H.; Liang, J.; Doberva, M.; Paranjape, K.; Lalancette, C.; Allegra, S.; Faucher, S. P.; Prévost, M. Local Adaptation of Legionella Pneumophila within a Hospital Hot Water System Increases Tolerance to Copper. Applied and Environmental Microbiology 2021, 87 (10). DOI:10.1128/aem.00242-21.

17. Cazals, M.; Bédard, E.; Faucher, S. P.; Prévost, M. Factors Affecting the Dynamics of Legionella Pneumophila, Nontuberculous Mycobacteria, and Their Host Vermamoeba Vermiformis in Premise Plumbing. ACS ES&T Water 2023, 3 (12), 3874–3883. DOI:10.1021/acsestwater.3c00288.

18. Martin, R. L.; Strom, O.; Song, Y.; Mena-Aguilar, D.; Rhoads, W. J.; Pruden, A.; Edwards, M. A. Copper Pipe, Lack of Corrosion Control, and Very Low pH May Have Influenced the Trajectory of the Flint Legionnaires’ Disease Outbreak. ACS ES&T Water 2022, 2 (8), 1440–1450. DOI:10.1021/acsestwater.2c00182.

19. Rhoads, W. J.; Keane, T.; Spencer, M. S.; Pruden, A.; Edwards, M. A. Did Municipal Water Distribution System Deficiencies Contribute to a Legionnaires’ Disease Outbreak in Quincy, IL? Environmental Science & Technology Letters 2020, 7 (12), 896–902. DOI:10.1021/acs.estlett.0c00637.

20. Cazals, M.; Bédard, E.; Doberva, M.; Faucher, S.; Prévost, M. Compromised Effectiveness of Thermal Inactivation of Legionella Pneumophila in Water Heater Sediments and Water, and Influence of the Presence of Vermamoeba Vermiformis. Microorganisms 2022, 10 (2), 443. DOI:10.3390/microorganisms10020443.

21. Nescerecka, A.; Hammes, F.; Juhna, T. A Pipeline for Developing and Testing Staining Protocols for Flow Cytometry, Demonstrated with SYBR Green I and Propidium Iodide Viability Staining. Journal of Microbiological Methods 2016, 131, 172–180. DOI:10.1016/j.mimet.2016.10.022.

22. Singh, A.; Yelverton, C. J.; Barnard, T. G. Rapid Quantification of the Total Viable Bacterial Population on Human Hands Using Flow Cytometry with SYBR® Green I. Cytometry Part B: Clinical Cytometry 2019, 96 (5), 397–403. DOI:10.1002/cyto.b.21776.

23. Wang, H.; Edwards, M.; Falkinham, J. O.; Pruden, A. Molecular Survey of the Occurrence of Legionella Spp., Mycobacterium Spp., Pseudomonas Aeruginosa, and Amoeba Hosts in Two Chloraminated Drinking Water Distribution Systems. Applied and Environmental Microbiology 2012, 78 (17). DOI:10.1128/aem.01492-12.

24. Rački, N.; Morisset, D.; Gutierrez-Aguirre, I.; Ravnikar, M. One-Step RT-Droplet Digital PCR: A Breakthrough in the Quantification of Waterborne RNA Viruses. Analytical and Bioanalytical Chemistry 2013, 406 (3), 661–667. DOI:10.1007/s00216-013-7476-y.

25. Callahan, B. J.; McMurdie, P. J.; Rosen, M. J.; Han, A. W.; Johnson, A. J.; Holmes, S. P. DADA2: High-Resolution Sample Inference from Illumina Amplicon Data. Nature Methods 2016, 13 (7), 581–583. DOI:10.1038/nmeth.3869.

26. R Core Team. R: A Language and Environment for Statistical Computing; R Foundation for Statistical Computing: Vienna, Austria, 2021. https://www.R-project.org/. (accessed 2025-03-10).

27. Callahan, B. RDP Taxonomic Training Data Formatted for DADA2 (RDP Trainset 18/Release 11.5). Zenodo (CERN European Organization for Nuclear Research) 2020. DOI: 10.5281/zenodo.4310151.

28. Schloss, P. D. Rarefaction Is Currently the Best Approach to Control for Uneven Sequencing Effort in Amplicon Sequence Analyses. mSphere 2024, 9 (2). DOI:10.1128/msphere.00354-23.

29. Oksanen, J.; Guillaume Blanchet, F.; Friendly, M.; Kindt, R.; Legendre, P.; McGlinn, D.; Minchin, P. R.; O’Hara, R. B.; Simpson, G. L.; Solymos, P.; Stevens, M. H. H.; Szoecs, E.; Wagner, H. vegan: Community Ecology Package; R package version 2.6-4, 2022. https://CRAN.R-project.org/package=vegan (accessed 2025-03-01).

30. Bates, D.; Maechler, M.; Bolker, B.; Walker, S. Fitting Linear Mixed-Effects Models Using lme4. J. Stat. Softw. 2015, 67 (1), 1–48. DOI: 10.18637/jss.v067.i01

31. Hothorn, T.; Bretz, F.; Westfall, P. Simultaneous Inference in General Parametric Models. Biometrical J. 2008, 50 (3), 346–363. DOI: 10.1002/bimj.200810425.

32. Love, M. I.; Huber, W.; Anders, S. Moderated Estimation of Fold Change and Dispersion for RNA-Seq Data with DESeq2. Genome Biol. 2014, 15 (12), 550. DOI: 10.1186/s13059-014-0550-8

33. van der Kooij, D.; Veenendaal, H. R.; Scheffer, W. J. H. Biofilm Formation and Multiplication of Legionella in a Model Warm Water System with Pipes of Copper, Stainless Steel and Cross-Linked Polyethylene. Water Research 2005, 39 (13), 2789–2798. DOI:10.1016/j.watres.2005.04.075.

34. Edwards, M.; Pruden, A.; Falkinham, J. O., III; Brazeau, R.; Williams, K.; Wang, H.; Martin, A.; Rhoads, W. Relationship Between Biodegradable Organic Matter and Pathogen Concentrations in Premise Plumbing. The Water Research Foundation, 2013. https://www.waterrf.org/research/projects/relationship-between-biodegradable-organic-matter-and-pathogen-concentrations

35. Martin, R. L.; Harrison, K.; Proctor, C. R.; Martin, A.; Williams, K.; Pruden, A.; Edwards, M. A. Chlorine Disinfection of Legionella Spp., L. Pneumophila, and Acanthamoeba under Warm Water Premise Plumbing Conditions. Microorganisms 2020, 8 (9), 1452. DOI:10.3390/microorganisms8091452.

36. Roman, F. A., Jr.; Byrne, T.; Martin, R. L.; Mena-Aguilar, D.; Smeltz, R. E.; Finkelstein, R.; Pruden, A.; Edwards, M. A. Retrospective Analysis of Drinking Water Microcosm Microbiomes Reveals an Apparent Antagonistic Relationship between *Neochlamydia* and *Legionella*. Environmental Science & Technology Letters. 2025, DOI: 10.1021/acs.estlett.5c00590

37. Ishida, K.; Sekizuka, T.; Hayashida, K.; Matsuo, J.; Takeuchi, F.; Kuroda, M.; Nakamura, S.; Yamazaki, T.; Yoshida, M.; Takahashi, K.; Nagai, H.; Sugimoto, C.; Yamaguchi, H. Amoebal Endosymbiont Neochlamydia Genome Sequence Illuminates the Bacterial Role in the Defense of the Host Amoebae against Legionella Pneumophila. PLOS ONE 2014. DOI:10.1371/journal.pone.0095166.

38. Goss, P. J.; Peccoud, J. Quantitative Modeling of Stochastic Systems in Molecular Biology by Using Stochastic Petri Nets. Proceedings of the National Academy of Sciences 1998, 95 (12), 6750–6755. DOI:10.1073/pnas.95.12.6750.

39. De Vrieze, J.; De Mulder, T.; Matassa, S.; Zhou, J.; Angenent, L. T.; Boon, N.; Verstraete, W. Stochasticity in Microbiology: Managing Unpredictability to Reach the Sustainable Development Goals. Microbial Biotechnology 2020, 13 (4), 829–843. DOI:10.1111/1751-7915.13575.

40. Zhou, J.; Ning, D. Stochastic Community Assembly: Does It Matter in Microbial Ecology? Microbiology and Molecular Biology Reviews 2017, 81 (4). DOI:10.1128/mmbr.00002-17.

