## Supporting Information (SI Tables and Figures) for "Influence of Copper Dose on *Mycobacterium avium* and *Legionella pneumophila* Growth in Premise Plumbing"

for

**Contents:**

**Supporting Information Table 1**: ddPCR assays

**Supporting Information Table 2**: Statistical Analysis of Total Cell Counts

**Supporting Information Table 3:** Statistical Analysis of *M. Avium* data

**Supporting Information Figure 1:** *Legionella pneumophila* and Total Cell Counts in the microcosms at the end of the acclimation phase

**Supporting Information Figure 2:** Total Cell Counts (TCC) in 2000 µg/L Cu condition over copper dosing period

**Supporting Information Figure 3:** Comparison between Legiolert™ and ddPCR targeting *L. pneumophila (Lp)*

**Supporting Information Figure 4:** Legiolert™ data collected across study

**Supporting Information Table 1.** Summary of ddPCR assays.

| Target | Supermix | Primers/Probes | Annealing Temperature (°C) | Reference |
|---|---|---|---|---|
| *L. pneumophila* | Probe | Forward: AAAGGCATGCAAGACGCTATG | 58.9 | (Nazarian et al., 2008)[1] |
| | | Reverse: GAAACTTGTTAAGAACGTCTTTCATTTG | | |
| | | Probe: FAM-TGGCGCTCAATTGGCTTTAACCGA | | |
| *M. avium* | Evagreen | Forward: AGAGTTTGATCCTGGCTCAG | 59.4 | (Wilton et al., 1992)[2] |
| | | Reverse: ACCAGAAGACATGCGTCTTG | | |

Thermal cycling for *L. pneumophila* amplification consisted of an initial enzyme activation at 95 °C for 10 minutes, followed by 40 cycles of denaturation at 94 °C for 30 seconds and annealing at 58.9 °C for 1 minute. This was followed by a final enzyme deactivation step at 98 °C for 10 minutes, with reactions held at 4 °C thereafter.

Thermal cycling for *M. avium* amplification consisted of an initial enzyme activation at 95 °C for 10 minutes, followed by 40 cycles of denaturation at 95°C for 30 seconds and annealing at 58.9 °C for 1 minute. This was followed by signal stabilization at 4°C for 5 minutes, then at 90 °C for an additional 5 minutes. Reactions were then held at 4°C thereafter.

To evaluate potential PCR inhibition, each sample was initially tested at three concentrations: undiluted, 1:10, and 1:100. Due to low *L. pneumophila* concentrations within the microcosms, undiluted samples were used when targeting *Lp*. In contrast, *M. avium* was typically quantified using 1:10 dilutions.

**Supporting Information Table 2.** Tukey contrasts on a linear mixed-effect model of total cell count by copper dosage on the final month of the copper dosing phase, over three sequential water changes, controlling for repeated microcosm measurements.

| Copper | 0 | 4 | 30 | 250 | 2000 |
|---|---|---|---|---|---|
| 0 | 1 | 0.980 | 1 | <0.001 | <0.001 |
| 4 | 0.980 | 1 | 0.978 | <0.001 | <0.001 |
| 30 | 1 | 0.978 | 1 | <0.001 | <0.001 |
| 250 | <0.001 | <0.001 | <0.001 | 1 | 0.411 |
| 2000 | <0.001 | <0.001 | <0.001 | 0.411 | 1 |

**Supporting Information Table 3.** Tukey contrasts on a linear mixed-effect model of *M. avium* ddPCR gene copies/mL by copper on the final month of the copper dosing phase, over three sequential water changes, controlling for repeated microcosm measurements.

| Copper | 0 | 4 | 30 | 250 | 2000 |
|---|---|---|---|---|---|
| 0 | 1 | 0.998 | 1 | <0.001 | 0.426 |
| 4 | 0.998 | 1 | 1 | <0.001 | 0.251 |
| 30 | 1 | 1 | 1 | <0.001 | 0.349 |
| 250 | <0.001 | <0.001 | <0.001 | 1 | 0.026 |
| 2000 | 0.426 | 0.251 | 0.349 | 0.026 | 1 |

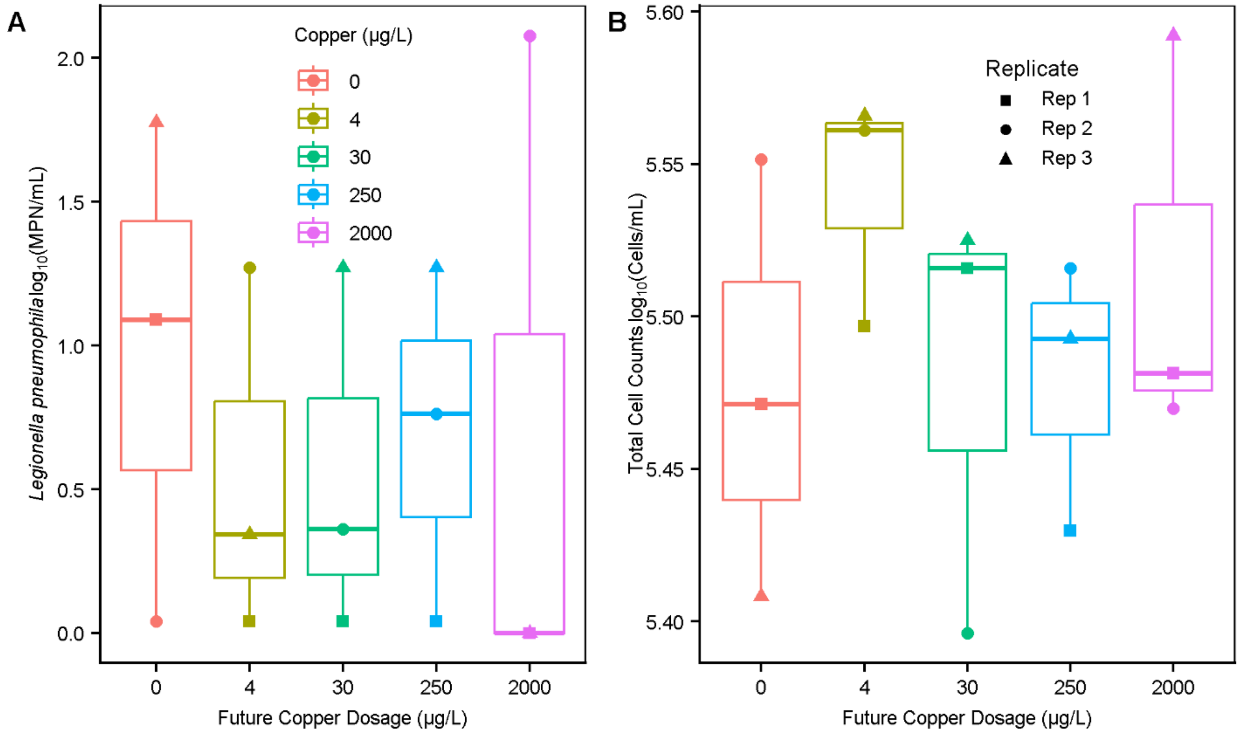


**Supporting Information Figure 1.** (A) Log *Lp* and (B) log TCC counts at the end of the acclimation phase, just prior to commencing copper dosing, grouped by the copper dosing they were destined to receive and with the shape corresponding to their replicate number. The data consists of 5 groups (that would later correspond to copper levels received) × 3 microcosms × 1 sampling event = 15 *Lp* and 15 TCC data points).

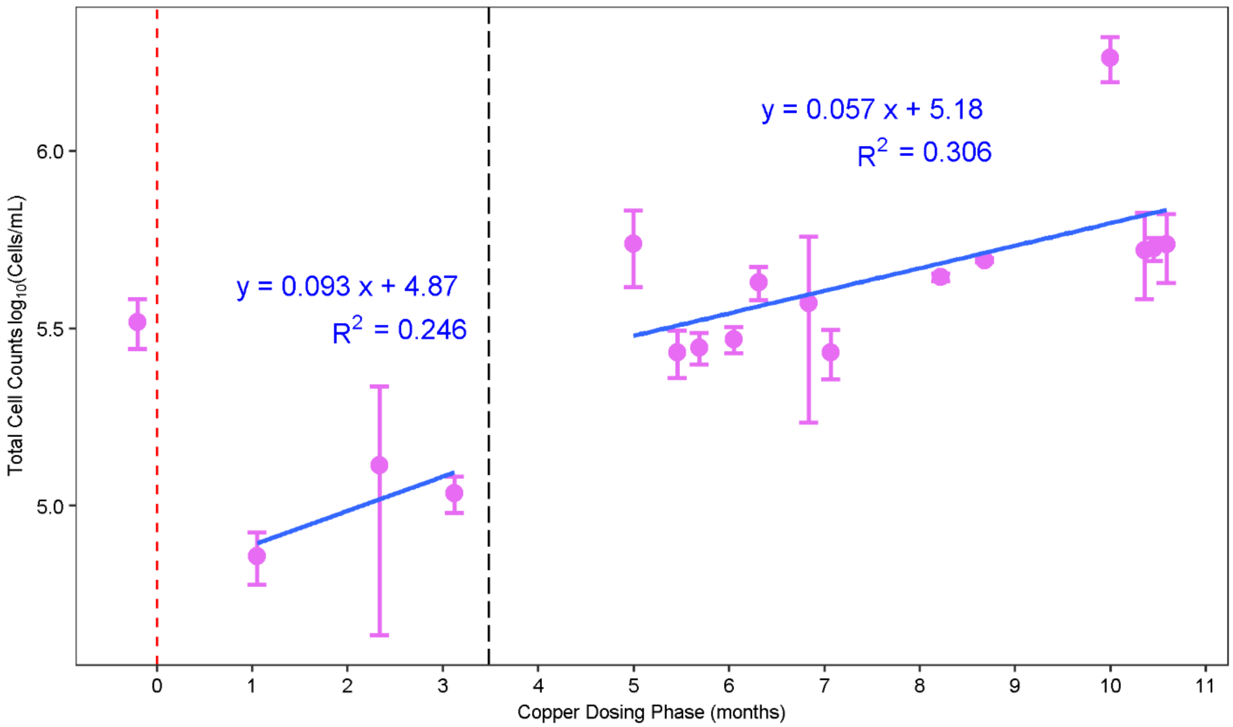


**Supporting Information Figure 2.** Mean ± standard deviation total cell counts of the 2000 µg/L microcosms throughout the copper-dosing phase (operated in biological triplicate n=3). The dashed red vertical line at Time 0 represents the beginning of copper dosing. The dashed black vertical line at Time ~3.5 represents the point at which all reactors were reinoculated with *Lp*.

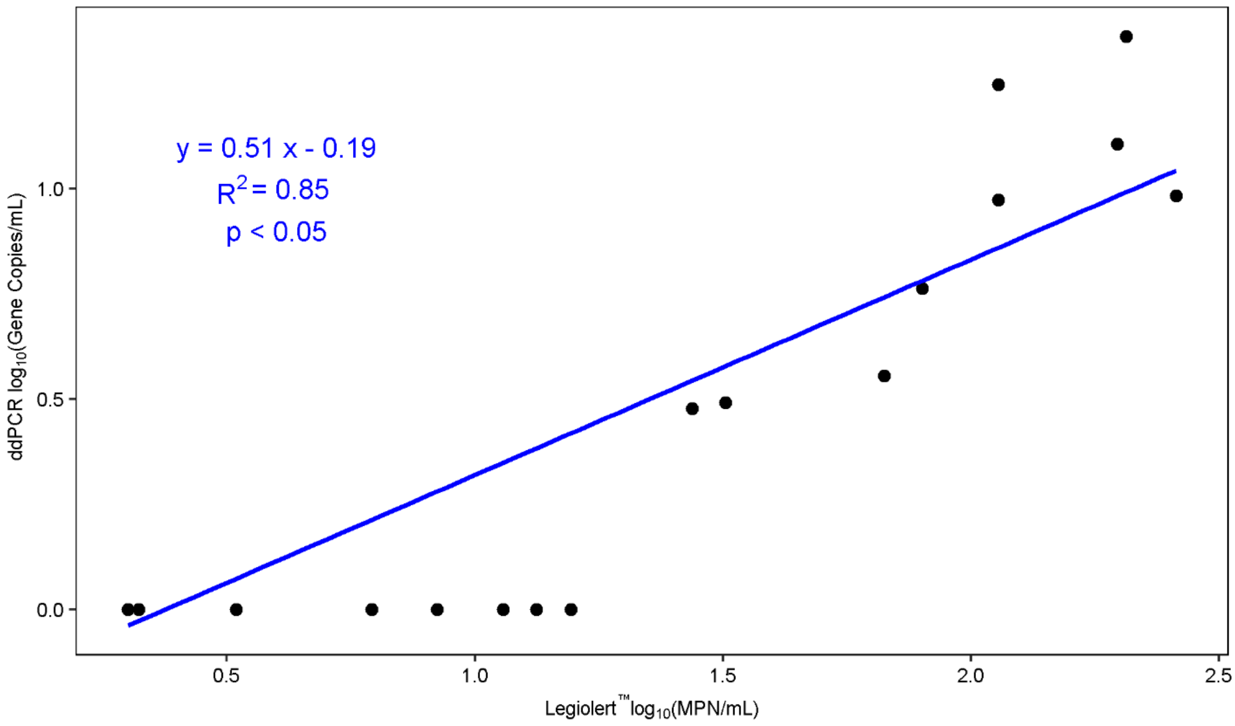


**Supporting Information Figure 3.** Comparison of Legiolert™ measurements as log(MPN/mL) to ddPCR measurements log(gene copies/mL) of *Lp* among samples for which both measurements were conducted. The blue line and equation represent the linear regression between the two variables. Data points plotted at a concentration of zero (gc/mL or MPN/mL) represent non-detects. The data consists of 5 copper levels × 3 microcosms (per copper level) × 3 sampling events = 45 data points.

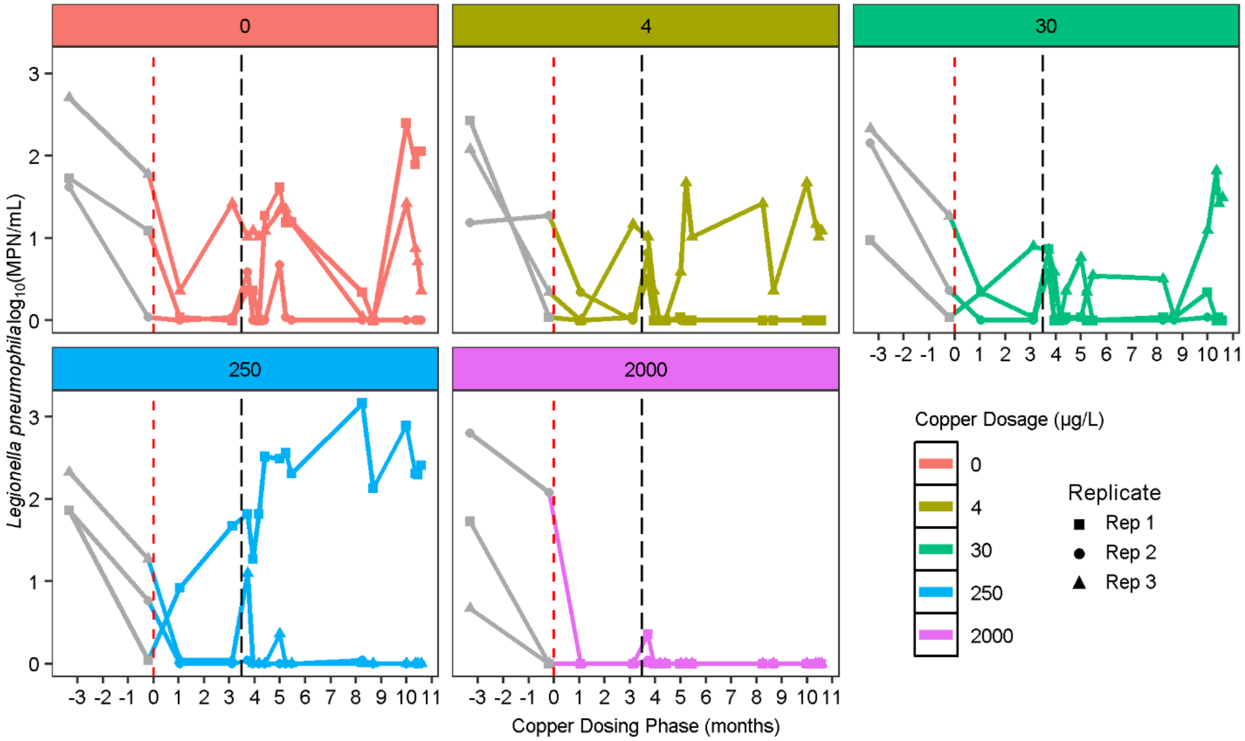


**Supporting Information Figure 4.** *Lp* quantification in MPN/mL determined by Legiolert™ tests for the three replicate microcosms dosed with copper over 11 months. The dashed red vertical line at Time 0 represents the time copper dosing began. The dashed black vertical line at ~ 3.5 months represents the point at which all microcosms were reinoculated with water from the original lab system containing *Lp* from which the microcosms had originally been seeded. The detection limit was 1 MPN/mL. For *Lp*, points plotted at an MPN/mL of zero indicate non-detects.

**References**


1. Nazarian, E. J.; Bopp, D. J.; Saylors, A.; Limberger, R. J.; Musser, K. A. Design and Implementation of a Protocol for the Detection of Legionella in Clinical and Environmental Samples. Diagnostic Microbiology and Infectious Disease **2008**, 62 (2), 125–132. DOI:10.1016/j.diagmicrobio.2008.05.004.

2. Wilton, S.; Cousins, D. Detection and Identification of Multiple Mycobacterial Pathogens by DNA Amplification in a Single Tube. Genome Research **1992**, 1 (4), 269–273. DOI:10.1101/gr.1.4.269.
